# Pro-Inflammatory Molecules Implicated in Multiple Sclerosis Divert the Development of Human Oligodendrocyte Lineage Cells

**DOI:** 10.1101/2024.12.17.628958

**Authors:** Gabriela J. Blaszczyk, Abdulshakour Mohammadnia, Valerio E. C. Piscopo, Julien Sirois, Qiao-Ling Cui, Moein Yaqubi, Thomas M. Durcan, Raphael Schneider, Jack P. Antel

## Abstract

**Background and Objectives:** Oligodendrocytes (OL) and their myelin-forming processes are targeted and lost during the disease course of Multiple Sclerosis (MS), targeted by infiltrating leukocytes and their effector cytokines. Myelin repair is considered to be dependent on recruitment and differentiation of oligodendrocyte precursor cells (OPCs). The basis of failure of re-myelination during the disease course of MS remains to be defined. The aim of this study is to determine the impact of pro-inflammatory molecules tumor necrosis factor ⍰ (TNF⍰) and interferon gamma (IFN_*γ*_) on the differentiation of human OPCs.

**Methods:** We generated human OPCs from induced pluripotent stem cells with a reporter gene under the OL-specific transcription factor SOX10. We treated the cells *in vitro* with TNF⍰ or IFN_*γ*_ and evaluated effects in terms of cell viability, expression of OL-lineage markers, and co-expression of astrocyte markers. To relate our findings to the molecular properties of OPCs as found in the MS brain we re-analyzed publicly available single nuclear RNAseq datasets.

**Results:** Our analysis indicated that both TNF⍰ and IFN_*γ*_ decreased the proportion of cells differentiating into the OL-lineage; consistent with previous reports. We now observe the TNF⍰ effect is linked to aberrant OPC differentiation. A subset of O4+, reporter-positive cells co-expressed the astrocytic marker Aquaporin-4 (AQP4). On the transcriptomic level, the cells acquire an astrocyte-like signature alongside a conserved reactive phenotype. Analysis of single-nuclear RNAseq datasets from human MS brain revealed a subset of OPCs expressing an astrocytic signature.

**Discussion:** In the context of MS, these results imply that OPCs are present but inhibited from differentiating along the OL-lineage, with a subset acquiring a reactive and stem-cell like phenotype, reducing their capacity to contribute towards repair. These findings help define a potential basis for the impaired myelin repair in MS and provide a prospective route for regenerative treatment.

## 1. Introduction

Multiple Sclerosis (MS) is a neuroinflammatory disease of the central nervous system (CNS), characterized by multi-focal demyelinating lesions underlying disease relapses. Extent of neurologic recovery and subsequent development of a progressive disease course are linked at least in part to extent of tissue repair (1), and has yet to be addressed with currently available therapeutics. Initial lesion formation reflects injury of oligodendrocytes (OLs) and their myelin membranes by direct contact with CNS-infiltrating immune cells (2) and their effector molecules(3). Progressive disease is linked with ongoing loss of OLs. Myelin repair mechanisms are dependent on recruitment and differentiation of OL progenitor cells (OPCs) shown to be present in the CNS parenchyma. Block of OPC differentiation ability has been observed in situ (1, 4), however the exact cause remains unknown.

Soluble mediators of tissue injury released by CNS-infiltrating leukocytes include pro-inflammatory molecules such as tumour necrosis factor-alpha (TNF⍰) and interferon-gamma (IFN_*γ*_). These have been shown to directly affect myelinating OLs as well as activate other resident glial cells (5). Both TNF⍰ and IFN_*γ*_ have been found to be cytotoxic to rodent derived primary OPCs in vitro (6-8). However, functional studies of primary human OPCs are limited by access as such cells arise only in later second trimester of development (9). Previous studies using human induced pluripotent stem cells (iPSCs) generated by genetic reprogramming (10) have shown that TNF⍰ and IFN_*γ*_ inhibited the terminal differentiation of OL lineage cells. These cytokines reduced process extension from primary mature human OLs in vitro without de- differentiation (11).

Aside from their canonical roles in generating myelinating OLs, OPCs can mediate a number of immune functions. In addition to their motility, it has been noted that OPCs have phagocytic capacity (12), release pro-inflammatory cytokines (13), and express major histocompatibility complex (MHC) molecules (14, 15) which are involved with leukocyte activation and recruitment. The term “reactive” OPC was coined by Simon et al. (2011) (16), where OPCs upregulated Neural/Glial antigen 2 (NG2, *CSPG4* gene) and increased proliferation in response to tissue injury. Koupourtidou et al. (2024) (17) contributed to these observations in which these reactive OPCs upregulated the same set of genes as astrocytes and microglia following stab wound injury, hinting at a conserved immune response amongst glial cells.

In the current study we used a growth factor-based protocol for deriving human OL lineage cells from iPSCs (18), including the use of a reporter gene under the OL-specific transcription factor SOX10. This approach allowed us to assess the effects of TNF⍰ and IFN_*γ*_ on early differentiation stages and lineage commitment of these cells. We observed that TNF⍰ exposure induces the upregulation of Aquaporin-4 (AQP4) *in vitro*. Bulk RNA sequencing (RNAseq) analysis of reporter positive OPCs indicated upregulation of astrocytic genes alongside an acquired reactive OPC signature following TNF⍰ exposure. Furthermore, reanalysis of previously published MS single nuclear RNAseq (snRNAseq) datasets confirmed the presence of *AQP4*+ OPCs *in situ*, preferentially localizing to active lesions in the brain.

## 2. Methods

### Standard Protocol Approvals, Registrations, and Patient Consents

The use of iPSCs and stem cells in this research was approved by the McGill University Health Centre Research Ethics Board (DURCAN_IPSC/2019–5374).

### Human iPSC culture and OPC differentiation

Previously characterized healthy control iPSC lines (3450, 3450-*SOX10*^*mO*^) generated in-house at the Early Drug Discovery Unit were used for this study (18, 19). iPSC-derived OL lineage cells were differentiated according to a recently published protocol (18). Cells were treated with TNF⍰ (100 ng/mL, ThermoFisher Scientific, Mississauga, ON) or IFN_*γ*_ (100 ng/mL, ThermoFisher Scientific, Mississauga, ON) for 4 to 6 days prior to fixation (days 18-21 of differentiation).

### Immunofluorescence Staining and Analysis

Cell viability (propodium iodide, PI, Invitrogen, Waltham, MA) and differentiation (O4, R&D systems, Oakville, ON) were determined by immunofluorescence as previously described in-detail (20). Briefly, cells were stained live with anti-O4 (1:200) and PI (1:200) for 15 minutes at 37 degrees. Post-fixation, cells were stained corresponding secondary antibodies (1:400) conjugated with Alexa Fluor 647 (Thermo Fisher Scientific, Missisauga ON). Plates were imaged with a 10X objective using a Zeiss Axio Observer fluorescence microscope (Carl Zeiss Canada, Toronto, ON) or the ImageXpress (Molecular Devices, San Jose, CA) high-content imaging platform following staining for O4 and PI. Cells were counted by a blinded individual using the ImageJ software or automatically using the ImageXpress software.

### Fluorescence Activated Cell Sorting and Flow Cytometry

For the iPSC-reporter line, sorting was based on reporter intensity using a previously described protocol (18), with the FACS Aria Fusion cell sorter (BD Biosciences, San Jose, CA). For phenotyping, cells were stained as previously described, using the LIVE/DEAD cell viability kit (Invitrogen, Waltham MA) and pre-conjugated antibodies (O4-APC, Miltenyi, Auburn, CA; AQP4-AF488, Bioss USA, Woburn, MA). Samples were acquired on the Attune flow cytometer (Thermo-Fisher Scientific, Mississauga, ON). Data was analyzed using the FlowJo software (BD Biosciences, San Jose, CA).

### Bulk RNAseq preparation and analysis

FACS-sorted 3450-*SOX10*^*mO*^ cells were collected, following treatment with TNF⍰ for four days. RNA was extracted using the Norgen (Thorold, ON) single cell RNA purification kit.

Library preparation, sequencing, quality check, alignment, quantification of raw read counts, and normalization of read counts were performed using the same methods as described in Mohammadnia et al.(21). As these samples did not exhibit significant heterogeneity, we used DESeq2 (22) for differential gene expression analysis following the methodology outlined in Pernin et al.(23). Significantly differentially expressed genes were identified using a threshold of log2 fold change > 1 and adjusted p-value cut-off of < 0.05. Hierarchical clustering was performed using Seaborn’s “clustermap” function (Citation: Waskom 2021, Journal of Open Source Software) with an average linkage method and correlation-based distances on log 2 transformed normalized read counts. We applied custom colormaps to highlight expression differences across conditions with row normalization and heatmaps were generated with matplotlib (Citation: Hunter 2007, Computing in Science and Engineering) and Seaborn to visualize.

Using the gseapy package, we performed single sample Gene Set Enrichment Analysis (ssGSEA) on the normalized read counts (Citation: Fang et al., 2023, Bioinformatics). The ssGSEA was run with rank-based normalization, and gene sets were defined from a GMT file of biological pathways (c5.go.bp.v2024.1.Hs.symbols.gmt).

All single-cell RNA-seq analyses were performed using the same pipeline employed in our previous publication (23) on the same Jakel et al.(24) and Absinta et al. (25) data sets.

### Statistical analysis

Statistical analyses were performed using Excel or GraphPad Prism software. Cell culture studies were performed with at least three individual replicates per experiment (see figure legends for details). Student’s t-test was used for comparisons between two groups or a Dunnett’s multiple comparisons test was used. P-values <0.05 were considered statistically significant.

### Data Availability

Sequencing data will be provided upon acceptance/publication of the manuscript.

## 3. Results

### Pro-inflammatory molecules do not induce cytotoxicity in human OPCs, but decrease the proportion of cells in the OL-lineage

We generated OL lineage cells from human iPSCs as described in Piscopo et al (2024), utilizing our previously characterized iPSC reporter line expressing the fluorophore mOrange under the control of the OL-specific transcription factor SOX10 (*SOX10*^*mO*^). We sorted *SOX10*^*mO*^ cells as previously described (18) (Figure S1), where the fluorescence intensity of the reporter is directly proportional to the developmental status of OL-lineage cells. Under basal conditions, (Figure 1A) around 45% of cells are mOrange medium, whereas 17% are mOrange high, indicating a majority of early OL-lineage cells. To study the effects of the selected molecules on our human OPCs, we simultaneously treated cells with our selected pro-inflammatory molecules while inducing differentiation. Exposure of the cells to TNF⍰ and IFN_*γ*_ resulted in a 50% reduction in the proportion of O4+ cells (TNF⍰ p = 0.0355, IFN_*γ*_ p= 0.0040). (Figure 1B). We used our previously published (20) glucose deprivation condition (NG) as a positive control for detection of cell death (p = 0.0007). In comparison, we did not observe significant cell death following exposure to TNF⍰ or IFN_*γ*_ (Figure 1C). This then prompted us to assess whether another glial cell type was compensating for this decrease in the proportion of OPCs, given how this protocol generates mixed-glial cultures (18). Interestingly, we observed that the percentage of AQP4+ astrocytes increased (p < 0.0001) following treatment with TNF⍰ (Figure 1D). Building on these findings, we next sought to elucidate the source of the increased proportion of AQP4+ in culture.

**Figure 1.**
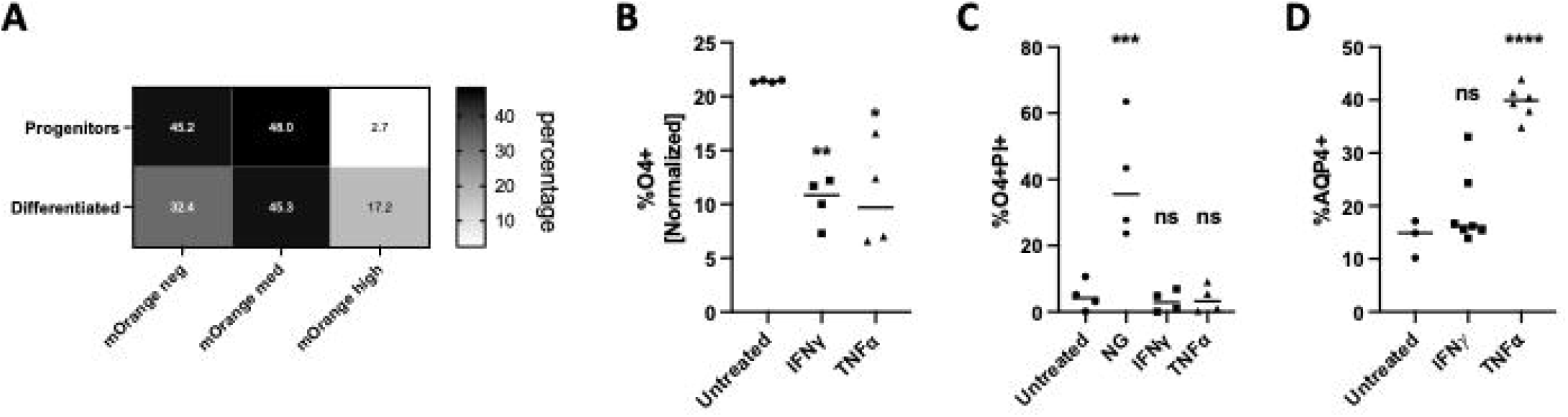
Pro-inflammatory molecules block human OPC differentiation. **(A)** iPSC-OPCs generated from our SOX10^mO^ reporter line, subset based on fluorescence intensity before and after 21-day differentiation, showing an increase in SOX10^mO^ with not all cells being terminally mature (mOrange high). Median percentage value of n=4 passages displayed in heat map. **(B)** Percentage of O4+ OPCs (normalized) as counted following immunofluorescence staining and treatment for 4 days. n = 4 passages. * p < 0.05, ** p < 0.01. **(C)** Proportion of O4+PI+ dead OPCs as measured by immunofluorescence staining following treatment for 4 days. n = 4 passages. *** p < 0.001. NG = no glucose, positive control. **(D)** Proportion of AQP4+ astrocytes in culture as measured by flow cytometry following treatment for 4-6 days, n = 3 passages. **** p < 0.0001.

### Increase in astrocyte proportions under inflammatory conditions *in vitro* derived from an initial OPC population

It has been suggested that OPCs may retain or acquire multilineage potential under certain conditions (26-28), although this has not yet been demonstrated in human cultures. For example, rodent OPCs exposed to fetal bovine serum have been shown to generate astrocytes (29). With fate mapping techniques in murine models, studies have shown that in both normal development (28) and stress conditions (30) OPCs are directed into astrocytes. To explore this possibility, we analyzed the O4 + population in our iPSC-derived cell lines for co-expression of other surface markers via flow cytometry. In response to cytokine exposure (Figures 2A-B), a proportion of the cells co-expressed the astrocyte marker AQP4 with TNF⍰ (p = 0.0046) treatment but not with IFN_*γ*_ (p = 0.7678). To establish that this population originated from the OL-lineage, we used our *SOX10*^*mO*^ reporter line (18). We observed that the AQP4+ O4+ sub-population of cells expressed *SOX10*^*mO*^ (Figure 2C), suggestive of an OL-lineage origin. These results indicate that a proportion of astrocytes in our cultures following TNF⍰ exposure originated from our *SOX10*^*mO*^ reporter positive OPCs.

**Figure 2.**
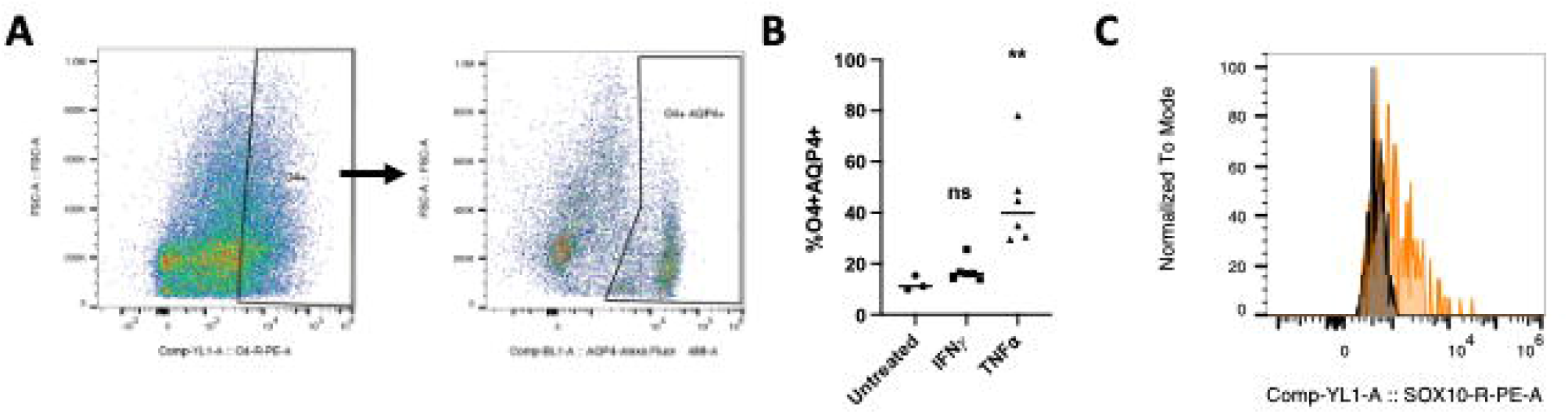
TNF⍰increases the proportion of OPCs co-expressing AQP4. **(A)** Representative gating scheme of OPC marker O4 (PE) co-expressing astrocyte marker AQP4 (AF488) in a TNF⍰ treated sample **(B)** Percentage of O4+ OPCs also expressing AQP4 as measured by flow cytometry. n = 3, treatment between 4-6 days, ** p < 0.01. **(C)** Histogram of SOX10^mO^ fluorescence intensity of O4+AQP4+ cells (orange) against a non-reporter control (black). Fluoresence intensity values normalized to mode.

### SOX10^mO^ OPCs acquire a reactive and astrocyte-like gene signature

To determine if the observed phenotypic responses of our TNF⍰ treated cells could also be seen at the transcriptomic level, we performed bulk RNA sequencing following cell sorting of our reporter line (18). The majority of O4+AQP4+ presented with an intermediate *SOX10*^*mO*^ intensity based on previously published criteria (18) (Figure S2). Following sequencing of our *SOX10*^*mO*^- sorted cells, principal component analysis revealed that the primary source of variance is reflected in PC1 (Figure 3A). TNF⍰ treatment led to changes in the gene expression profile in comparison with the control group (Figure 3B). Our *SOX10*^*mO*^ cells had an enrichment for terms related to immune processes in response to TNF⍰, as well as terms for glial cell proliferation and activation (Figure 3C). A previous study (17) observed a common glial response to injury amongst microglia, astrocytes and OPCs in response to a stab wound injury. To verify if this is true in our model, we performed a single sample gene set enrichment analysis (ssGSEA) of this conserved gene signature and found a majority (87%) of genes upregulated (Figure 3D). We observed a significant enrichment score of neural progenitor cell (NPC) proliferation signature (Figure 4A), further contributing to the notion of our TNF⍰ treated OPCs exhibiting the previously described reactive OPC phenotype (16). To confirm our *in vitro* findings of OPCs acquiring astrocytic characteristics, we compared our TNF⍰ treated OPCs to OL differentiation and astrocyte gene signatures (Figures 4B-C). We observed downregulation of mature OL genes (*MBP, PLP1*) and OPC identity (*PDGFRA, GPR17*) (Figure 4B). Furthermore, we observed a significant enrichment score of astrocytic differentiation (Figure 4C) and upregulation of key genes, confirming our flow cytometry findings. This suggests that in addition to a reactive OPC gene signature, OPCs exposed to TNF⍰ acquire gene signatures associated with NPC proliferation and astrocytic differentiation, highlighting a complex response to inflammation.

**Figure 3:**
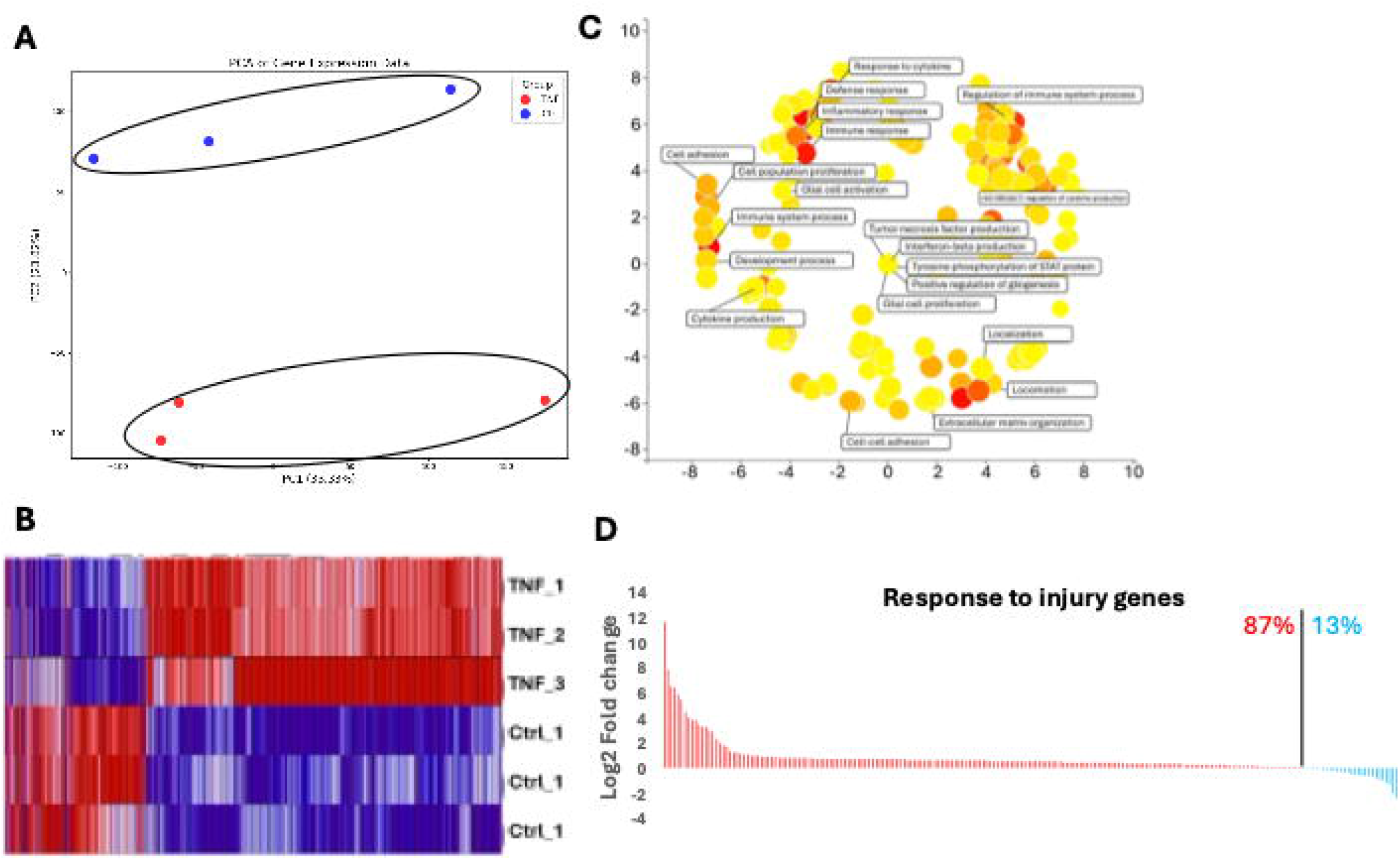
Transcriptome changes in iPSC-derived SOX10-med cells upon treatment with TNF⍰. **(A)** Principal Component Analysis (PCA) using all detected genes reveals a clear effect of TNF treatment in PC2. Control samples are shown in blue, and treated samples in red. Normalized read counts from all detected genes were used. **(B)** Hierarchical clustering was performed on significantly differentially regulated genes (adjusted p-value < 0.05 and log2 fold change > 1). Normalized read counts were used for clustering, and row normalization was applied to visualize the heatmap. **(C)** Gene Ontology (GO) analysis of significantly upregulated genes was conducted using g:Profiler, and GO terms were summarized and visualized with the Revigo tool. **(D)** Genes related to injury response in glial cells as published by Kopourditou et al (2024) were more enriched in TNF-treated cells compared to control cells. Genes are visualized using log2 fold change following single-sample gene set enrichment analysis.

**Figure 4:**
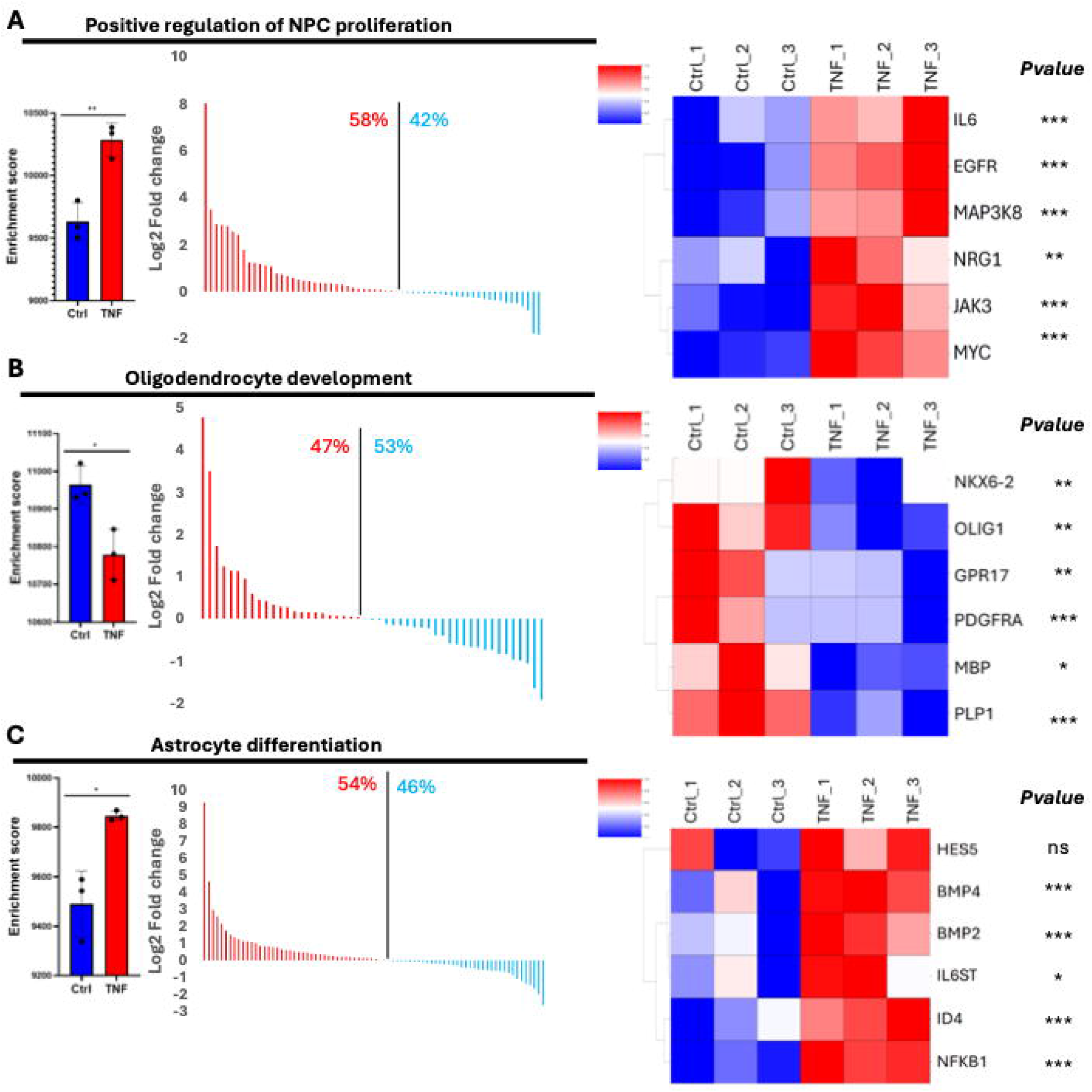
TNF⍰ treatment of iPSC-derived SOX10-med cells deviates glial differentiation program and includes a neural progenitor cell (NPC) proliferation signature. **(A)** Single-sample gene set enrichment analysis (ssGSEA) of the NPC proliferation signature shows significant enrichment in TNF-treated cells compared to control cells (left panel). Log2 fold changes indicate a higher number of genes with increased expression in this signature upon TNF treatment (middle panel). Examples of significantly upregulated NPC proliferation genes are visualized in a heatmap, with hierarchical clustering performed using normalized read counts, and DESeq2-calculated p-values provided for each gene (right panel). **(B)** ssGSEA of oligodendrocyte development genes using normalized read counts shows reduced enrichment of this signature in TNF-treated cells (left panel). Log2 fold change analysis reveals a higher proportion of genes with elevated expression in control cells compared to TNF-treated cells (middle panel). **(C)** ssGSEA of astrocyte differentiation genes indicates greater enrichment of this signature under TNF treatment conditions (left panel) and a higher proportion of genes with increased expression in this signature (middle panel). Examples of significantly upregulated genes involved in astrocyte differentiation are shown in the heatmap. Statistical significance is denoted as follows: ns = not significant, * = p < 0.01, ** = p < 0.001, *** = p < 0.0001 (DESeq2 results).

### OPCs in the MS brain exhibit an astrocyte-like gene signature

To address the hypothesis that a mechanism altering the balance of resident OPC populations in favour of astrocytes could limit repair in the MS brain, we reanalyzed publicly available single nuclear RNAseq datasets (24, 25). We investigated the OPC populations marked by *PDGFRA* and *CSPG4* expression (Figures 5A-B). In this population of OPCs, 5% co-expressed *AQP4* (Figure 5C). Comparative analysis of *AQP4*+ OPCs to top 100 genes of an astrocytic gene signature revealed that, *AQP4*+ OPCs are upregulating genes that include *SOX9* and *GFAP* (Figure 5D). We found this subpopulation to preferentially localize to the core of an MS lesion (Figure S3). The identification of this OPC population with an astrocytic gene signature localized within human MS brain lesions may prove a novel mechanism preventing adequate remyelination in an inflammatory context.

**Figure 5:**
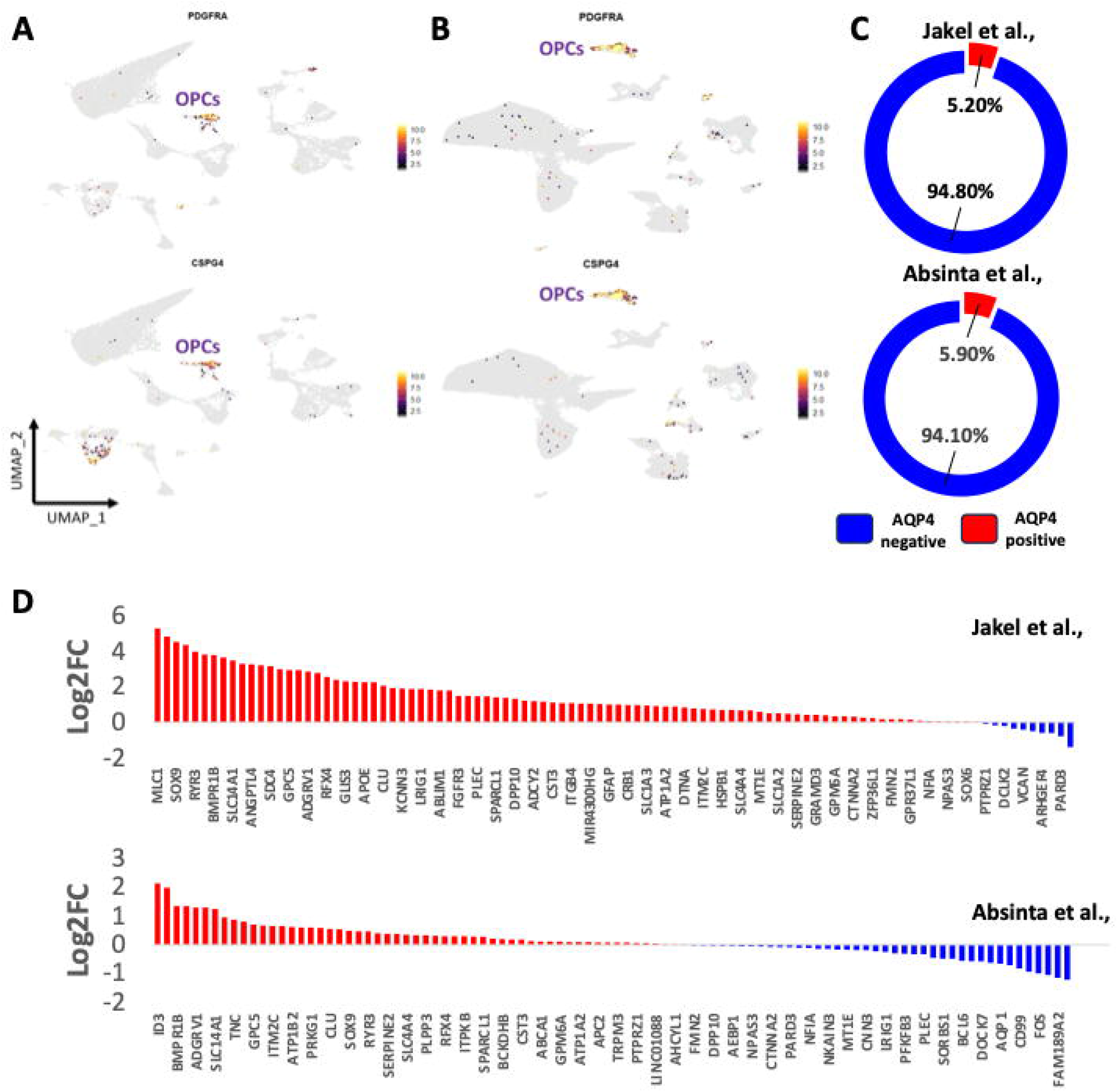
Identification and characterization of oligodendrocyte progenitor cells (OPCs) with astrocytic signatures in human single-nucleus RNA sequencing (snRNA-seq) datasets. **(A-B)** Analysis, clustering, and annotation of human snRNA-seq datasets derived from the studies by Jakel et al. **(A)** and Absinta et al. **(B).** OPC populations were identified using specific marker genes, notably PDGFRA and CSPG4/NG2. **(C)** A subset of OPCs was found to express AQP4, a marker typically associated with astrocytes. AQP4-positive cells are indicated in red, whereas cells lacking AQP4 expression are depicted in blue. **(D)** Comparative analysis of AQP4- positive OPCs versus a gene signature based on top 100 genes in the Jakel et al. (2019) astrocyte cluster, with the log2 fold change (Log2FC) depicted in bar plots. Top panel presents data from Jakel et al., 2019, and bottom panel from Absinta et al., 2021, with upregulated genes shown in red and downregulated genes in blue.

## 6. Discussion

Upon CNS infiltration during MS, leukocytes and glial cells (astrocytes and microglia), will release cytokines which have a direct effect on OLs and surrounding glia (5, 31), contributing to an overall pro-inflammatory microenvironment. These cytokines, among others, are found to be increased in the spinal fluid and peripheral blood of people with MS (31-33). Microarray analysis of MS lesions has shown an increase in cytokine expression and downstream regulatory pathways (34). Furthermore, spinal fluid lymphocyte gene expression of both TNF⍰ and IFN_*γ*_ have been found to be correlated with lesion load as seen by MRI (32), highlighting the relevance of these cytokines to MS disease pathogenesis.

Maturity state is a significant determinant in the response of glial cells to inflammatory mediators. Moore at al. (35) observed TNF⍰ -mediated apoptosis of early A2B5+ glial precursor cells and PDGFR⍰ + OPCs isolated from 14- to 20- week fetal human brain samples. Cui, D’Abate et al (36) showed that the myelination potential of these cells was more limited when derived from tissue earlier in gestation. Pernin et al (37), observed that these mediators induced

process retraction in mature OLs rather than cytotoxicity. Our use of a growth-factor based iPSC model recapitulates early OL commitment and early remyelination. We found reduced OPCs (O4+) in response to TNF⍰ and IFN_*γ*_ individually without the induction of cytotoxicity (as seen in rodent models and early human glial precursors), alongside an increase in AQP4+ astrocytes (Figure 1). These findings demonstrate that pro-inflammatory contexts in our model may alter the balance of other glial populations, in parallel to reducing OPC differentiation rather than promoting OPC death.

It has been suggested that OPCs may retain the capacity to generate astrocytes in adulthood as a response to pathological insults (30, 38, 39). Following neurogenesis in the developing brain, gliogenesis begins with the generation of astrocytes followed by oligodendrocytes, as reviewed by Miller and Gauthier (40). Thus, early OL-lineage cells may retain the capacity for astrocytic generation. It has been shown that certain *in vitro* conditions may induce the developmental trajectory of OPCs towards an astrocytic fate (29). Several subsequent studies of spinal cord injury (30), brain ischemia (38), and glioma (39) have suggested that this is possible across other disease models as well. Here, we consider whether a similar phenomenon is being induced by pro-inflammatory molecules. Using our previously characterized iPSC reporter line (*SOX10*^*mO*^) and analyzing the expression of cell surface markers (O4, AQP4), we have observed an increase in an astroglial marker (AQP4) in our cultures following TNF⍰ exposure (Figure 1D), and an emergence of OPC subpopulation co-expressing AQP4 (Figure 2). In addition to reinforcing previous findings regarding the retention of multi-lineage potential in OPCs (26-28), this could unveil an alternative mechanism by which the endogenous OPC-mediated repair pathway in the CNS is compromised in MS conditions. Rather than differentiating into myelinating OLs, OPCs may instead give rise to astrocytes, contributing to the failure of proper myelin repair OPCs are immune competent, which is demonstrated by the expression of MHC molecules (15), for example. Rodent OPC exposure to IFN_*γ*_ as well as TNF⍰ has shown to increase phagocytic activity (41), and OPCs have been shown to engulf axons/synapses in the developing brain (12). A reactive OPC can be characterized by the increase of NG2 and proliferation (16). Simon et al.

(16) first noted this reactive OPC phenotype in rodents as a hallmark after brain injury. We further observed that gene signatures related to neural stem cells (NPC) and cell cycle were significantly upregulated (Figure 4A). *CSPG4* (NG2) was moderately upregulated although not significant (data not shown). However, the underlying mechanisms driving this response and its potential consequences for the broader OL population remain unclear. Our findings suggest that inflammatory stimuli can shift OPCs toward an immune-like function at the expense of their oligodendroglial precursor role, thereby potentially impairing their capacity for remyelination (Figures 3-4), expanding on previous observations in rodents (13). By acquiring stem like signatures and skewing the differentiation response of OPCs following injury (Figure 4), the balance of glial cells becomes affected with the increased generation of astrocytes as seen in our cultures (Figure 1).

This study demonstrates that early human OL-lineage cells retain multilineage potential alongside their reactive capacity, likely as an adaptive response to stressors such as the pro-inflammatory conditions characteristic of MS. We observed a decrease of differentiation (Figure 1), in line with previous studies in different models, alongside the emergence of a population identified by the surface markers AQP4, O4 and our reporter (Figure 2). Furthermore, we observed an enrichment in proliferative and astrocytic gene signatures (Figure 4), which may indicate a skewed differentiation response. Relating back to the MS brain, we observed a subpopulation of cells expressing *AQP4* within the OPC cluster (Figure 5) with an astrocytic gene signature, suggesting that this is a biological response with disease relevance. These findings highlight a potential therapeutic avenue—strategies aimed at promoting OPC differentiation along the OL lineage while inhibiting astrocytic conversion could enhance remyelination and mitigate disease progression.

## 5. Acknowledgements

We thank lab members Lama Fawaz, Vanessa Omana, Wolfgang Reintsch, Andrea Krahn, and Genevieve Dorval for technical or administrative assistance. The flow cytometry work and cell sorting were performed at The Neuro’s Early Drug Discovery Unit’s Flow Cytometry Core Facility.

## 6. Author Contributions

G.J.B, A.M., V.E.C.P., Q.L.C., and J.P.A contributed substantially to the conception and design of the study. G.J.B, A.M., V.E.C.P., and J.P.A drafted a significant portion of the manuscript or figures. G.J.B., A.M., V.E.C.P., J.S., Q.L.C., and M.Y. contributed to acquisition and analysis of data. T.M.D., R.S., and J.P.A. contributed to critical manuscript revision and oversaw the project. All authors approved the final version of the manuscript.

