## Supplemental Figures for "Pro-Inflammatory Molecules Implicated in Multiple Sclerosis Divert the Development of Human Oligodendrocyte Lineage Cells"

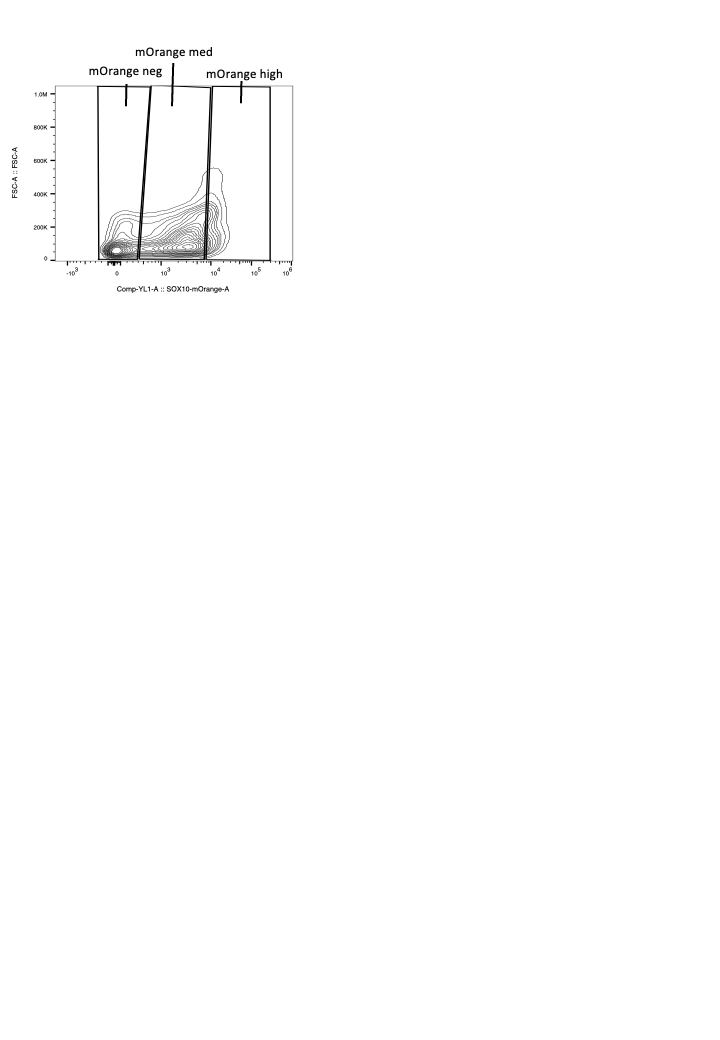
 **Supplementary Figures**

**Figure S1. Gating strategy for the SOX10^mO^ reporter line.**

Gating strategy where reporter intensity correlates with maturation stage along the oligodendroglial lineage. mOrange neg = no reporter expression. mOrange med = mild reporter intensity. mOrange high = high reporter intensity.

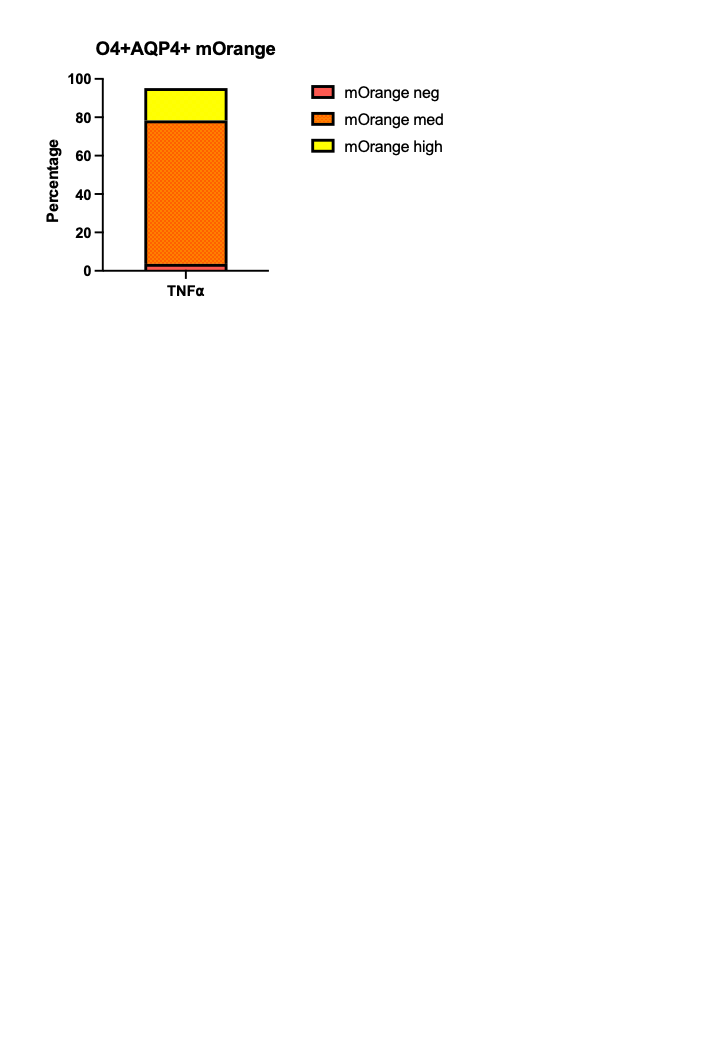
**Figure S2. SOX10^mO^ reporter intensity of O4+AQP4+ cells.**

A barplot representing gating of O4+AQP4+ cells on SOX10^mO^ reporter intensity as described in figure S1. A majority of O4+AQP4+ cells were found to be mOrange medium. Yellow indicates mOrange high cells, orange indicates mOrange medium cells, red indicates mOrange negative cells.

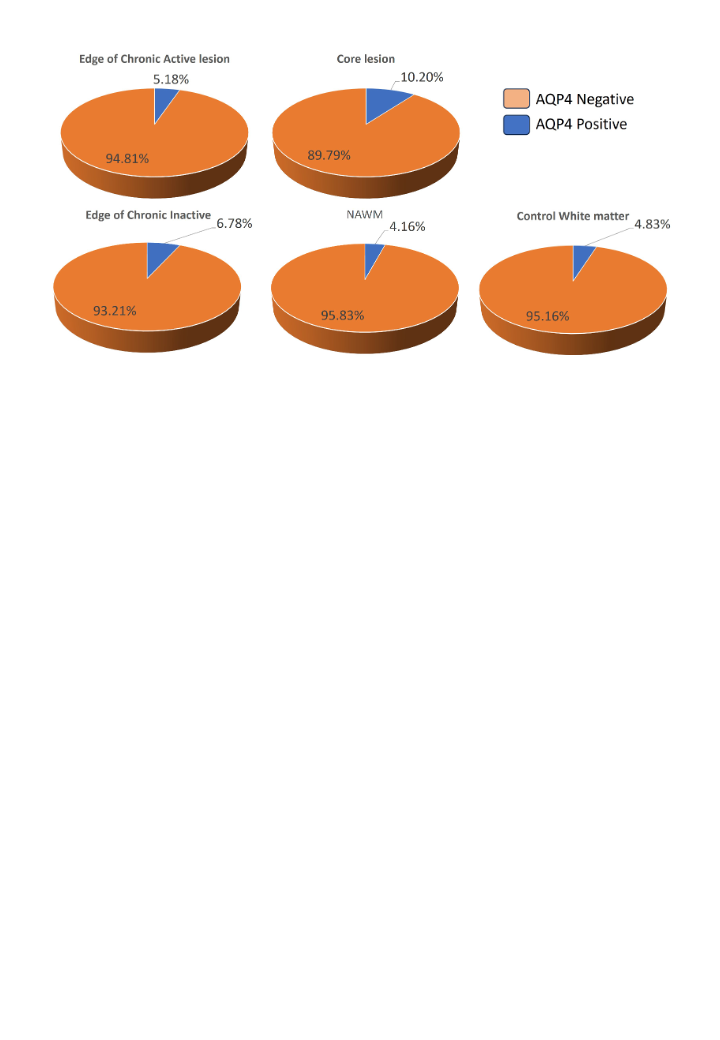
**Figure S3. Localization of AQP4+ OPCs in the MS brain.**

Pie plots representing percentage of AQP4+ cells within OPCs found in various regions of the MS brain, data-mined from the Absinta et al. single nuclear RNAseq (snRNAseq) dataset. Highest proportion of AQP4+ OPCs found to be localized in the core of an MS lesion. Orange represents AQP4- OPCs, blue represents AQP4+ OPCs.
